# Robust hydrogel-integrated microsystems enabled by enhanced interfacial bonding strength

**DOI:** 10.1101/2021.12.31.474677

**Authors:** Sang Wook Bae, Yong-Woo Kim, Jeong-Yun Sun, Sunghoon Kwon

## Abstract

Noncovalent hydrogels, compared to covalent hydrogels, have distinctive advantages including biocompatibility and self-healing property but tend to have poor mechanical robustness, thus restricting their application spectrum. A clue to increase utility of such soft hydrogels without chemical bulk modification can be witnessed in biological organ walls where soft mucous epithelial layers are juxtaposed with tough connective tissues. Perhaps, similarly, bonding noncovalent hydrogels to stronger materials, such as tough hydrogels, might be a viable approach for increasing stability and scalability as well as creating novel functions for hydrogel-based systems. However when attempting to bond these two materials, each of the four existing hydrogel-hydrogel bonding method has practical shortcomings. In this work, we introduce a mucosa-inspired bonding method that realizes interfacial bonding of noncovalent hydrogels to tough, hybrid hydrogels without external glue or bulk modification of the noncovalent gel while preserving interfacial micropatterns. The procedure is simple and we confirmed broad applicability with various noncovalent hydrogels and tough hydrogels. We demonstrated the utility of our bonding method with novel applications regarding in vitro assay, soft robotics and biologically inspired systems.

## Result

Hydrogel materials^1-5^ are now being adapted to fields including soft robotics^6,7^, and tissue engineering^8,9^. As a sub-category, naturally derived `noncovalent hydrogels` have good biocompatibility^10^ and self-healing property^11^. They are physically crosslinked hydrogels formed by weak bonds such as hydrogen bonds or ionic bonds. In case of agarose, although it tends to have poor mechanical properties, many biomimetic, fluidic systems integrated with these hydrogels were introduced due to their excellent biocompatibility. For example, microchannel patterned, or cell-laden agar/agarose gels were used in microsystems for cell culturing^12^, organism imaging^13^, point-of-care devices^14^, or ordered assembly^15^. Yet, applications of such purely noncovalent hydrogel was restricted since it tends to have low interfacial bonding strength with other polymers or solids^16^. To overcome this problem, it becomes important to develope methods that would enable strong adhesion between agarose hydrogel and solid. Previously introduced hydrogel-to-solid fixation strategies could be categoriezed into either mechanical clamping^17,18^ or chemical adhesion^19-22^. Mechanical clamping uses jigs to maintain contact between hydrogel and other solids without slippage or bursting. But they are temporary approaches and for non-trival hydrogel-solid adhesions such as forming kinetic joints^2,3^ or sealing channel-tube junctions^12^, it would be more effective to use chemical adhesion. Chemical adhesion approaches use external glues^19,20^ and/or incorporate solid-adhering functional groups into the hydrogel’s polymer backbone^2,3,21,22^. These recent advances in chemical hydrogel-solid adhesion techniques opened new opportunities for applying many hydrogels into microsystem designing. Concordantly, to develope new systems robustly integrated with noncovalent hydrogels, such as agarose, it would be desirable to develope a new chemical adhesion method that enables strong adhesion between solids and noncovalent hydrogels while preserving the bulk property of the hydrogel.

In this work, we introduce a facile method for increasing interfacial bonding strength of agarose hydrogel against solids via an interface-toughening hydrogel, designated as `hybrid hydrogel` (**Figure 1**). In other words, this hybrid hydrogel acts as a two-sided tape between agarose hydrogel and solids. Although we concentrated our analysis mostly on utilizing agarose hydrogel, our hybrid-hydrogel tape showed wider applicability to other noncovalent hydrogels as well. The bonding method requires no mechanical clamps and no bulk modification of the agarose hydrogel’s polymer backbone. It is also compatible with preserving micropatterns within the bonding interface, unlike liquid glues. Our hybrid hydrogel consists of a physically crosslinked polymer network (aldehyde-activated agarose) embedded with UV-crosslinkable monomers (acrylamide, AAm) and initiator (APS or Irgacure 2959). The mechanism of bonding agarose hydrogel with this hybrid hydrogel is two-fold. First is ‘network intertwining’. In detail, when agarose is placed on the hybrid hydrogel, this leads to passive diffusion of UV-crosslinkable monomers from the hybrid hydrogel into the noncovalent hydrogel matrix. After photo- or thermal polymerization, covalent network forms not only the second polymer network within the hybrid hydrogel but also the bonding interface that intertwines between agarose hydrogel and hybrid hydrogel. The interpenetrating network (IPN) formed at the two hydrogels` interface was observable with SEM and EDS **(Figure S1)**. Second part of the mechanism is `network crosslinking` by Schiff base^23-26^ formation. When this bond formed between the hybrid hydrogel’s aldehyde groups (**Figure S2**) and amine groups within the interfacial layer’s PAAm, the bonding strength was further increased compared to using ‘network intertwining’ alone. So due to the synergy of `network intertwining` and

**Figure 1.**
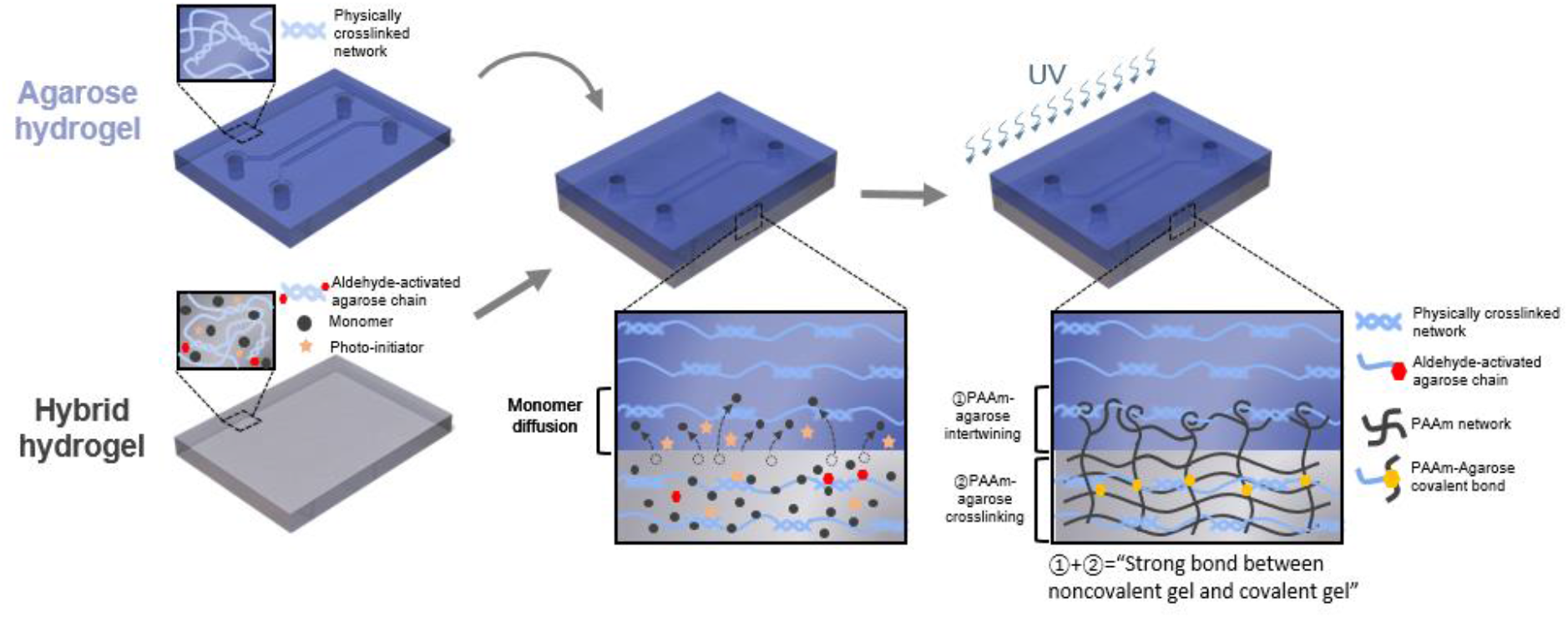
Schematic of bonding noncovalent agarose hydrogel to covalent hybrid hydrogel. Schematic of the overall bonding technique by combining ‘network intertwining’ and ‘network crosslinking’. A micropatterned agarose hydrogel is placed on a UV-reactive monomer-containing hybrid. Subsequently, monomer and photoinitiator diffuses into the agarose hydrogel followed by UV induced crosslinking which creates an intertwined network at the interface.

`network crosslinking` this single hybrid hydrogel was able to bond with various unmodified noncovalent hydrogels. The hybrid gel could also simultaneously covalently crosslink onto elastomers and nonporous solids on its opposite surface using previously introduced solid surface treatments^2,3^. We also provide evidence that our bonding mechanism may be applicable to other noncovalent hydrogels such as agar, gelatin, alginate, and chitosan. With this new bonding technique, we were able to fabricate novel noncovalent hydrogel-solid integrated structures and functionalities.

To measure the bonding strength between agarose gel and agarose-acrylamide(AAm) hybrid hydrogel^27-30^, we performed flatwise tensile test (**Figure 2a,b)**. We observed dose dependency of adhesive failure between agarose hydrogel and agarose:PAAm hybrid hydrogel containing 5∼15 w/v% AAm:MBA=19:1 concentration (**Figure 2c)**. Beyond 15 w/w%, cohesive failure occured due to the interface bonding strength exceeding the cohesive strength of agarose gel (**Figure S3a**). Additionally, we discovered that bonding strength could be further increased by adjusting the aldehyde-activation level of agarose chain within the hybrid hydrogel (**Figure 2d**). This aldehyde-activation was conducted by treating agarose powder with varying concentrations of oxidative sodium periodate (NaIO_4_)^24,26^. NMR spectrum of NaIO_4_-treated agarose showed, compared to normal agarose, a new peak at 209.5ppm which corresponds to aldehyde group as expected (**Figure S2b**,**c**). FT-IR analysis of hybrid hydrogels with varying ‘NaIO_4_ treatment duration’ showed that with increasing duration, the imine (C=N, 1565cm^−1^) peak of schiff-base increased while the amide II (1565cm^−1^) peak of PAAm chain decreased (**Figure 3a,b**). This indicated that PAAm amides were being consumed to form imine bond with aldehyde groups on the agarose chain, thus confirming our hypothesis. The force-extension curves of these experiments showed that adjusting AAm concentration within hybrid hydrogel changed the modulus of the construct (**Figure S3b)**, while adjusting aldehyde activation within the hybrid’s agarose network changed the stretchability of the construct (**Figure S3c)**. However, the level of agarose aldehyde activation had no significant effect on the fracture energy^31^ or stretchability of the hybrid hydrogel itself (**Figure S3d**).

**Figure 2.**
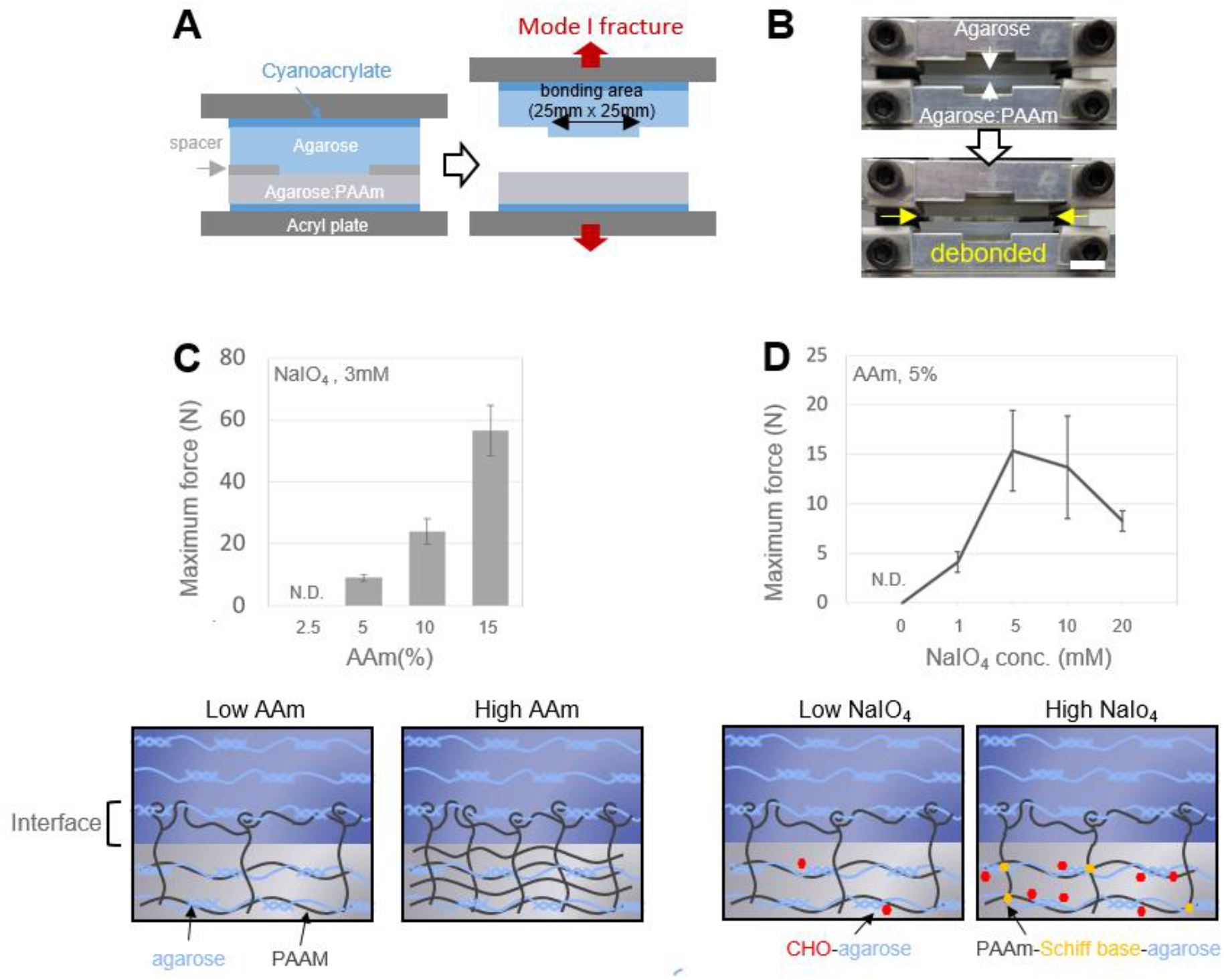
Validation of bonding mechanism. Bonding strength between agarose and hybrid hydrogel can be tuned by Acrylamide concentration and NaIO_4_ treatment concentration. **A, B**. Illustration(A) and demonstration (B) of debonding test based on a standard flatwise-tensile test (ASTM C297) test for measuring bonding force between agarose gel and agarose:PAAm hybrid gel. The bonded area was 25mm x 25mm. Cyanoacrylate was used for bonding hydrogels onto acryl plates. PET spacer or glass spacer was used to decrease the bonding area compared to the cyanoacrylate-bonded areas. **C, D**. Maximum force measured between tensile stress initiation and debonding between 3% agarose gel and various hybrid gels. Error bars are standard deviation of n=3∼4. N.D. indicates too low bonding strength to be measured.. **C**. Maximum forces measured with hybrid gels contained varying AAm monomer concentration. hybrid gels contained 3% agarose activated with 2mM NaIO4 for 6 hours, 10% AAm:MBA=19:1, 0.05 w/v% APS, and 0.15% TEMED. **D**. Maximum forces measured with hybrid gels made with agarose activated with varying sodium periodate concentration. Hybrid gels contained 3% agarose activated with sodium periodate for 3hours, 5%(w/v) AAm:MBA=19:1, 0.05% w/v% APS, 0.15% TEMED. Scale bar is 2.5mm (B).

**Figure 3.**
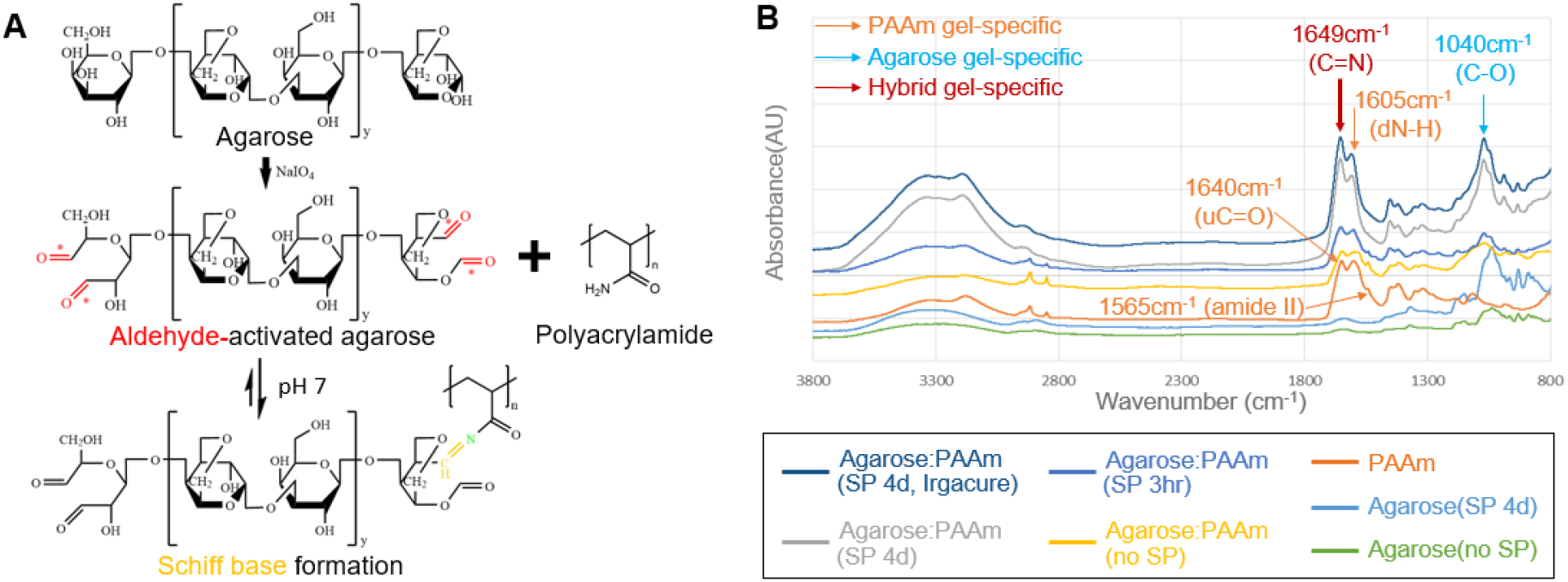
Validation of Schiff-base formation between aldehyde-agarose and polyacrylamide. **A**. Chemical structure of agarose, aldehyde-activated agarose, polyacrylamide, and formation of Schiff base. **B**. FT-IR absorbance spectra of various hydrogels. It is noticeable that only in hybrid hydrogels containing aldehyde-activated agarose, amide II peak (1565cm^−1^)^24^ of PAAm disappears and the peak(1649cm^−1^) increases which is indicative of imine(C=N)^35^ formation. The size variation of 2850cm^−1^ peak and 2920cm^−1^ peak from PAAm^35^ could be explained by difference in acrylamide monomer polymerization level^38^. SP indicates sodium periodate treatment of agarose (3hr: 3hours, 4d: 4 days). AAm crosslinking was initiated by APS except one sample (Agarose:PAAm, SP 4d, Irgacure 2959).

Additionally, changing the interfacial layer thickness had insignificant effect on bonding strength (**Figure S4a, b**). Based on our observation of the agarose:hybrid debonding pattern, this insensitivity of bonding strength to monomer diffusion time can be explained by agarose cohesive strength being stronger than interface-to-hybrid gel bonding strength in our experiment conditions (more detailed explanation is given in Figure S4d). For this experiment, the interfacial layer thickness was varied by changing the duration of monomer diffusion, and using Irgacure 2959 as AAm crosslinking initiator, and not APS because APS loses reactivity over time by oxygen exposure (**Figure S5**).

Based on these results, we applied this bonding method to bonding noncovlanet hydrogels with solid materials. First, we bonded noncovalent agarose gel with oxide-containing solid materials by using our hybrid hydrogel as a two-sided tape. We designed a 5 layer construct (glass-hybrid-agarose-hybrid-aluminum) that was bonded together by a single UV exposure step (**Figure 4a,b**). The resulting construct (bonding area=25mm x 30mm) was able to withstand a shear force of 3kgf (**Figure 4c-e)**. We were successful in producing higher bonding strength, compared to simple adhesion, between PDMS slab and other noncovalent hydrogels such as agar, alginate and gelatin using agarose:PAAm hybrid hydrogel as a two-sided tape (**Figure 4f, Figure S6**). Therefore, our noncovalent hydrogel-hybrid hydrogel bonding mechanism is not based on thermal or ionic gelation which works in bonding hydrogels with identical gelling mechanism (e.g. hydrogen bonding, ionic bonding). It also differs from previous chemical hydrogel-to-hydrogel bonding method where both hydrogel parts had to contain crosslinkable monomers^21^.

**Figure 4.**
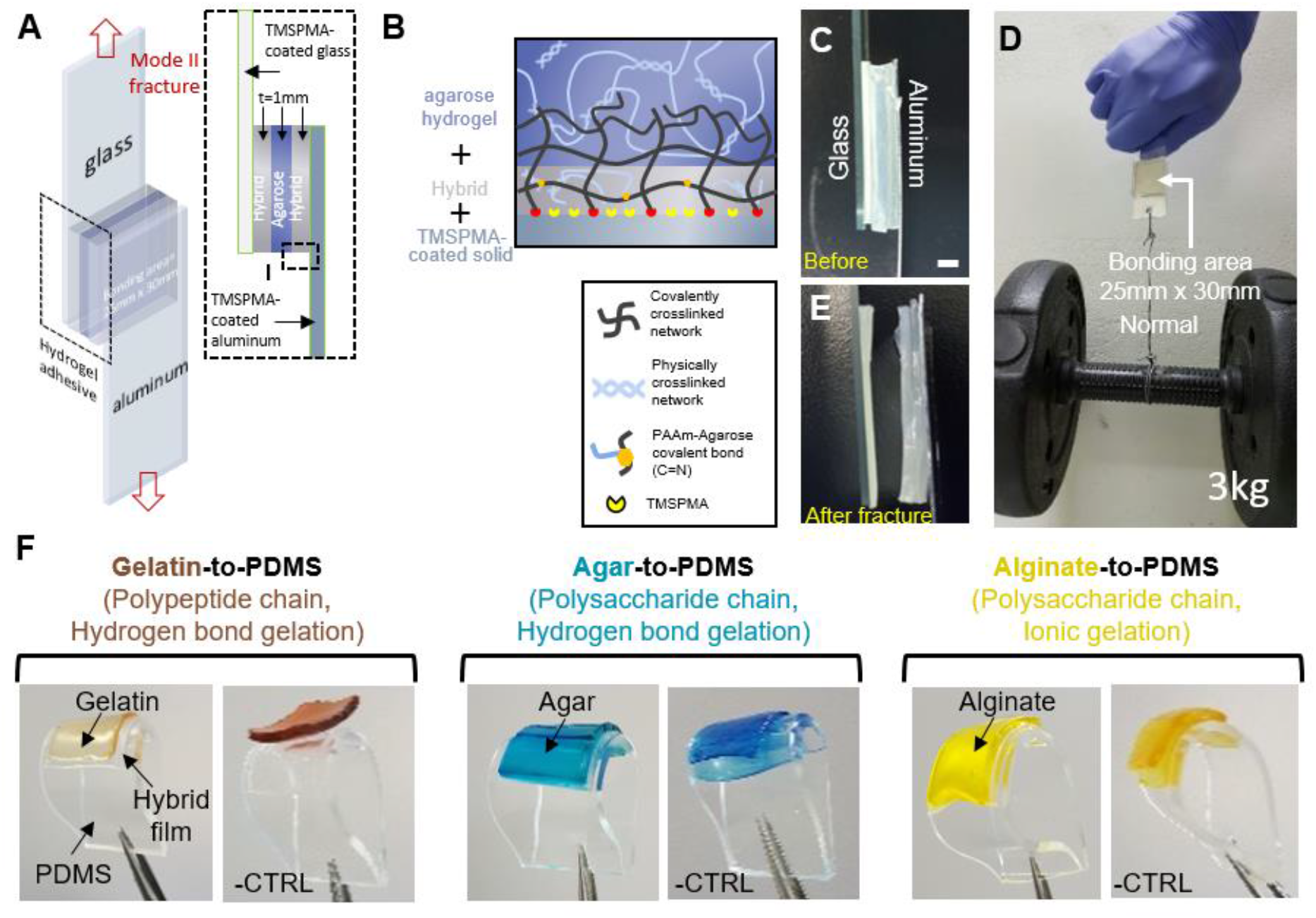
Expanding to bonding with solid materials and other noncovalent hydrogels. **A-E**. Schematic (A,B) and demonstration (C-E) of a 5 layer bonded sample. A 5 layer construct containing glass, aluminum, noncovalent agarose gel, and covalent Agarose:PAAm hybrid gel was made. This 5 layer construct of 25mm x 30mm bonding area was able to withstand 3kgf. **F**. bonding PDMS with agar, alginate, and gelatin using an agarose:PAAm hybrid gel film as a two-sided ape. Hydrogels were either UV-bonded with free monomer-containing hybrid hydrogel (left side images of each pair) or partially UV-cured hybrid hydrogel which lacks sufficient diffusible monomers (images indicated as -CTRL). Noncovalent hydrogels were pre-mixed with food dye for visualization. Scale bar is 1mm (C).

Previously, fabricating a robust agarose hydrogel-based microfluidic system required mechanical clamping between microchannel-patterned hydrogel and solid surface^17,18^, thus limiting scalability and design flexibility. But with our bonding scheme, we were able to robustly bond microchannel-patterned agarose gel onto glass substrates without mechanical clamping (**Figure 5a-e, Figure S7**). When fluid (Rhodamine B) was injected, the 250um x 100um cross-section channel withstood the maximum pressure generated by the syringe pump’s maximum flow rate (50 uL/sec, 2kPa). This was significantly higher than that obtained from a previously introduced agar-based microchannel system (0.22Pa) ^16^.

**Figure 5.**
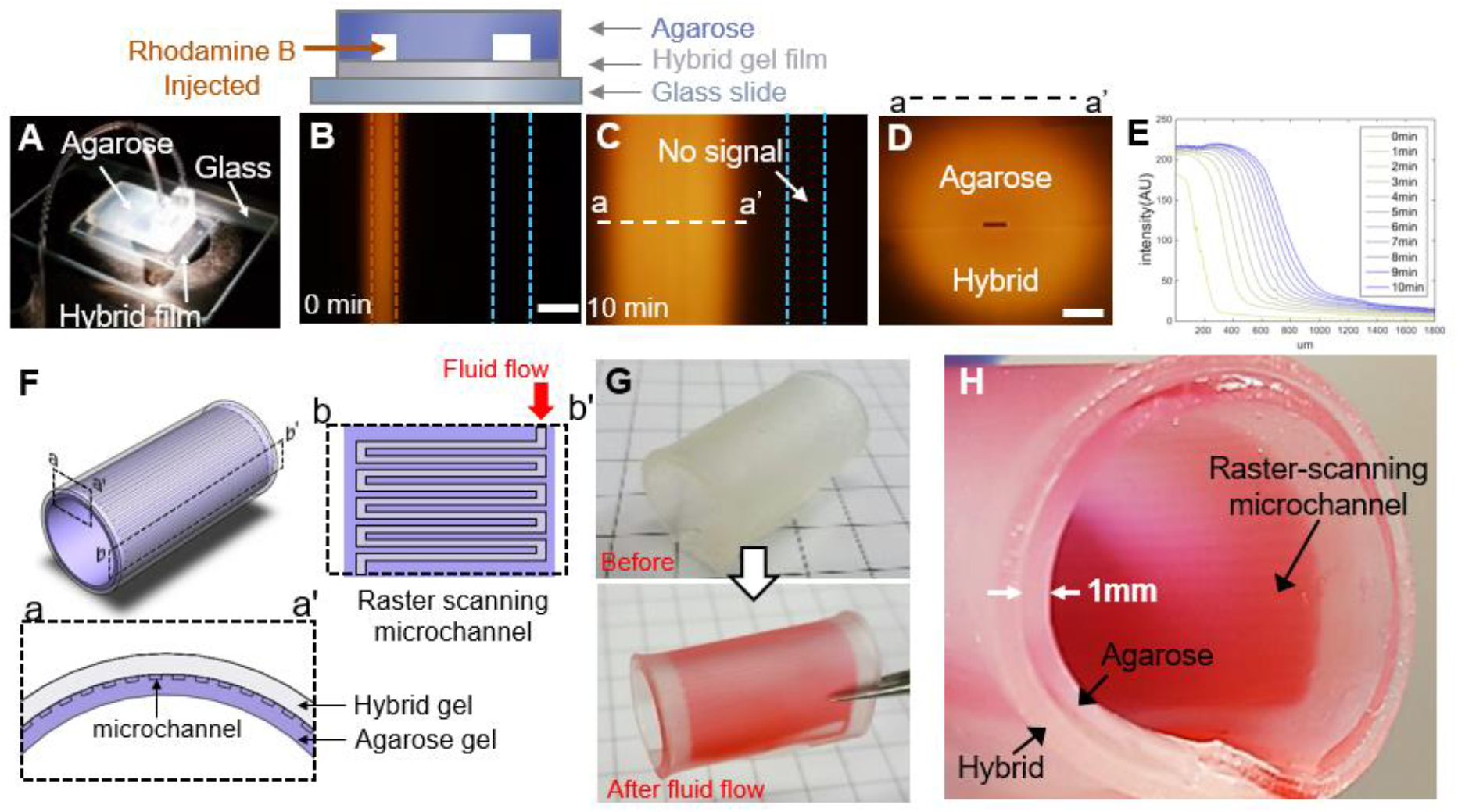
Validation of interfacial micro-patterning and application to organ model. **A-D**. A mechanical anchorage-free diffusive microfluidics demonstration. Rhodamine B was injected into a microchannel patterned between the agarose to hybrid bonding interface. After 10 minutes of dye injection, no signal was detected inside the closed channel (blue dashed line) on the right side of the dye-injected target channel (red dashed line), indicating successful channel sealing (C). Cross-section of the channel showed that dye diffused uniformly for all directions(D). E. Quantified time-lapse fluorescence diffusion profile. **F-H**. A hollow organ tissue-mimicking hydrogel system containing a raster scanning microchannel for fluid flow. Red food-dye was injected to demonstrate diffusive fluid flow (G,H). Scale bars are 500um (B,D). Grids in (G) are 10mm x 10mm.

Naturally derived noncovalent hydrogels are usually too fragile to mimic biological organs. Via our bonding approach, our hybrid hydrogel, which was tough against bending stress (**Figure S6**), could be used to structurally support fragile agarose hydrogel. As a demonstration, we fabricated an intestine-like cylindric hydrogel structure. Even though it contained thin (1mm) agarose **(Figure 5f-h)** or chitosan **(Figure S8)** hydrogel, the structure withstood such significant bending stress. Noticably, agar-based tough hydrogel^30^ (σ_fracture_ =0.2-0.7Mpa, ε_fracture_=180-260%) have similar mechanical property of small intestine^32^ (σ_fracture_=0.9Mpa, ε_fracture_=140%). The intermediate raster scanning microchannel enabled diffusive fluid flow which could be useful in preimplant applications for cell-enriching nutrient/biomolecule supply.

We also created a novel oligonucleotide-retrieving system by joining our agarose microfluidic structure with DNA electrophoresis (**Figure 6a-d**). Although inlets/outlets created by puncturing holes on hydrogels are, compared to stiff elastomers(e.g. PDMS), too fragile to tightly seal inserted tubes, our bonding method provided tight sealing between inserted glass tube and hydrogel channel inlet/outlet (**Figure 6d, Figure S9**). This enabled robust target DNA suction without fluid leakage or pressure loss. Our device was capable of selectively retrieving target DNA bands (1.4kbp, 1kbp, 500bp) from DNA ladder samples (**Figure 6e-g**). This is, to our knowledge, the first microfluidic target retrieval from electrophoretically size-separated sample. Our method is more straitforward compared to gel extraction method, which usually requires labor-intensive gel cutting, gel melting, sample purification, and strong chemicals (e.g. chaotropic salt) that damages oligonucleotide double helix (**Figure 6h, Figure S10**). We performed T7 endonuclease I assay which showed that our fluidic retrieval method retrieves target samples without double strand denaturing **(Figure 6i,j)**.

**Figure 6.**
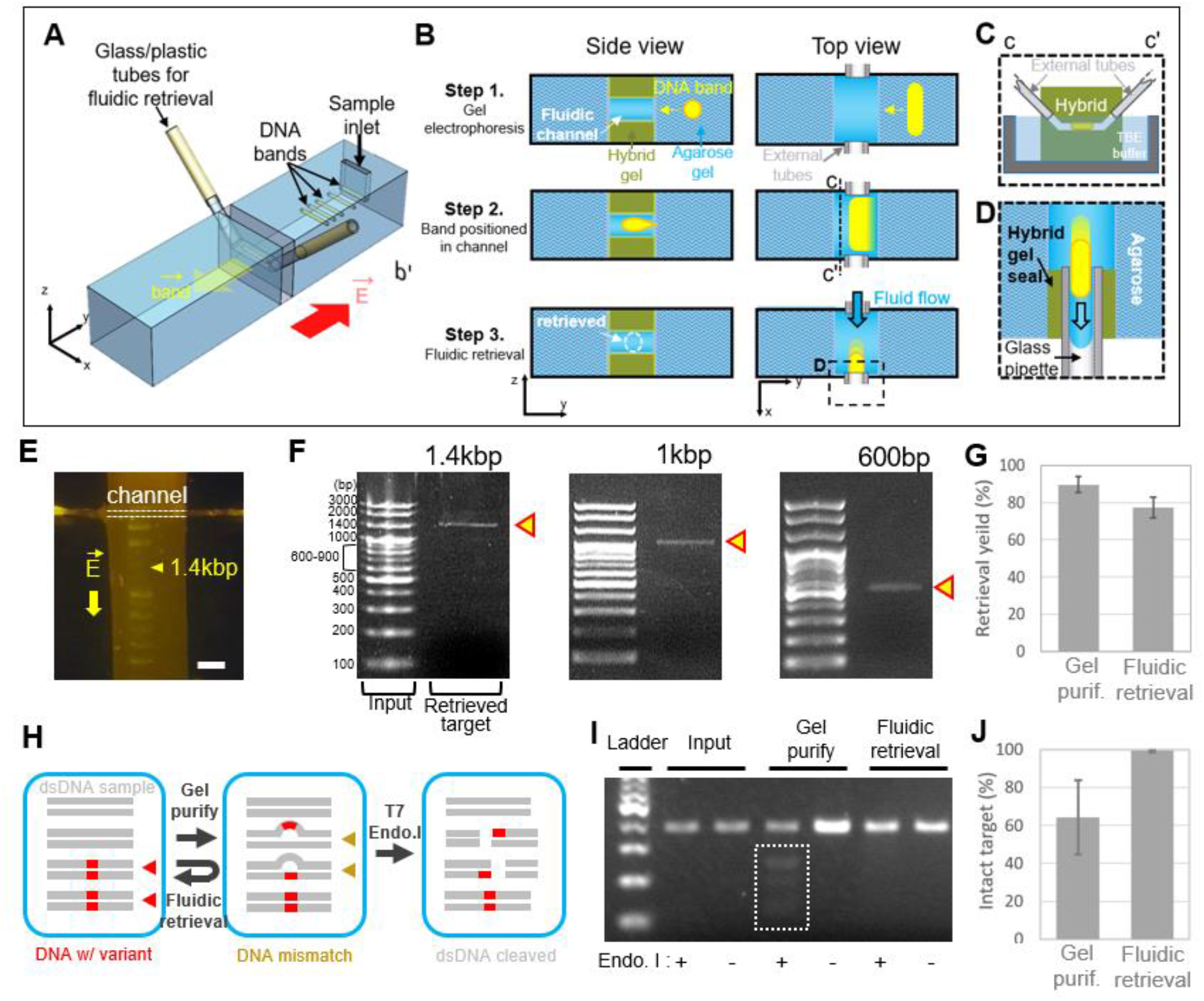
Hydrogel-based microfluidic electrophoresis sample retrieval system. **A-D**. Overall structure (A) and schematic (B-D) of a microfluidic DNA electrophoresis sample retriever. **E**,**F**. The target DNA sample (length 1.4kbp) was retrieved (E) and subsequently reloaded into a conventional agarose gel electrophoresis setup (F) to check purity. Experiment was repeated with 1kbp and 600bp targets. **G**. Comparison of 1.4kbp ladder retrieval yield between commercial gel purification kit (Qiagen) and fluidic retrieval method. (n=3) Quantity was measured by Qubit DNA broad-range kit (Invitrogen Inc.). **H**. Schematic showing the effect of retrieval method on double strand DNA (dsDNA) structure. Details are in figure S10. **I**. Example result of T7 endonuclease I treatment. Only gel-purified samples treated with T7 endonuclease I show cleaved fragments (dotted box). **J**. Percent of intact target (∼400bp) was measured as the proportion of intact target over total dsDNA content of the sample. (n=3). Scale bar is 5mm (E).

In summmary, we developed a agarose gel-to-solid bonding method that utilizes a hydrogel-formed two-sided tape. Our approach requires no mechanical anchorage, yet showed applicability to other noncovalent hydrogels without design modification of the hydrogel tape. The method also preserves interfacial micropatterns. With our approach, we increased stability and scalability of hydrogel-containing structures as well as creating various novel functionalities. We anticipate utilizing our approach to decipher further novel applications of noncovalent hydrogels in various bioengineering fields including in vitro assays, soft robotics, and tissue engineering.

### Experimental Section

#### Materials

Acrylamide (AAm, cat. no. A8887), *N,N*′-Methylenebis(acrylamide) (MBAA, cat. no. 146072), *N,N,N*′,*N*′-Tetramethylethylenediamine (TEMED, cat. no. T9281), Sodium alginate (cat. no. A2033), Gelatin (cat. no. G1890), Sodium tripolyphosphate (TPP, cat. no. 238503), Calcium sulphate dihydrate (cat. no. C3771), Benzophenone (cat. no. B9300), and Sodium periodate (cat. no. 311448) was purchased from Sigma Aldrich. Irgacure 2959 (cat. no. 55047962) was purchased from BASF. Ammonium persulfate (APS, cat. no. A2013), 1M Tris-HCl buffer (pH 7.4), and 19:1 AAm:MBAA (40 wt.%) solution was purchased from Biosesang (Korea). Agarose (cat. no. AG113-250) was purchased from Biofact. Agar (cat. no. 214010) was purchased from BD. Low melting agarose (LMA, cat. no. 32830) was purchased from Affymetrix. 50% Glutaraldehyde (cat. no. A10500) was purchased from Alfa Aesar. Cyanoacrylate (cat. no. 406) was purchased from Loctite. Aluminum plates (cat. no. FCK43) were purchased from Agami Modeling (Korea). Acryl plates were purchased from Acrylzip (Korea). 100bp ladder was purchased from Biosesang (cat. no. 15628-019). Ecoflex was purchased from Smooth-On^Inc^.

### Preparing hybrid hydrogel

Unless otherwise stated, % means w/w%. Agarose activation was conducted by first suspending agarose powder in DI water containing 1-20mM of sodium perdioate. The mixture was sealed to protect from light and stirred gently at room temperature for various duration (from 3 hours to several days). The resulting activated agarose powder was washed with DI water several times. For the agarose-AAm hybrid hydrogel, solution containing 5-20%(w/v) 19:1=Acrylamide(AAm): *N,N*′-Methylenebisacrylamide(MBAA) solution, 0.015 w/v% Ammonium persulphate (APS), was degassed, mixed with a microwave-heated 6% aldehyde-activated agarose (in DI water) solution in a ventilated hood. After the solution cooled to below 50°C, 0.15% *N,N,N*′,*N*′-Tetramethylethylenediamine (TEMED) was mixed. The solution was then poured into ice-cold glass mold, covered with ice-cold slide-glass, and put in a 4°C refrigerator for agarose gelation. When using Irgacure 2959, 0.2% Irgacure was used instead of APS and the TEMED adding step is omitted. For the alginate-AAm hybrid hydrogel, we referred to the previous publication^1^ with slight modification. Briefly, a 100mM Tris-HCl buffer-based solution containing 2% alginate, 12.05% AAm, 0.017% MBAA, 0.2% Irgacure 2959 (or 0.015 w/v% APS) was thoroughly degassed and quickly mixed with 20mM Calcium Sulphate (CaSO_4_), 2mM Sodium tripolyphosphate (TPP) slurry and poured onto a glass mold or hydrogel mold. When using APS instead of Irgacure, the slurrry additionally contained 0.25 v/v% TEMED. The pregel was let to physically crosslink in a vacuum chamber for 10-60 minutes.

#### Agarose hydrogel to agarose:AAm hydrogel bonding procedure

When using APS/TEMED based crosslinking, the bonding side of the noncovalent gel (1-3% agarose in DI or 0.5x TBE buffer) was treated with benzophenone (BP, 10 w/v% in absolute ethanol) for 1 minute, washed with absolute ethanol twice, and dried at atmospheric condition. Then the gel was transferred onto a hybrid hydrogel and exposed to UV irradiation for interfacial crosslinking (17mw/cm^2^ for 15-30 minutes). When using Irgacure 2959 based crosslinking, no BP treatment was needed.

#### Bonding strength measurement

We prepared agarose/hybrid gel construct having 2.5mm x 2.5mm bonding area. The hybrid gel side was bonded to an acrylic plate with cyanoacrylate. A custom jig was used to mount the acrylic plate-bonded gel construct at the bottom base of the tensile tester (MCT 1150, A&D Korea). We then coated the agarose side of the gel construct with cyanoacrylate. Immediately after this, another acrylic plate mounted on the upper crosshead of the tensile tester was let to contact the cyanoacrylate-coated agarose gel and bond. After 3-5 minutes of incubation and zero-point calibration, the tensile test was conducted at 10mm/min pulling speed until the agarose/hybrid gel interface fully de-bonded. We measured the ultimate tensile stress at the point of adhesive failure between agarose gel and hybrid gel.

### Fracture test

We measured the fracture energy of the hybrid hydrogel (containing 5% AAm:MBAA=19:1, 0-200mM NaIO_4_) with a previously introduced method^20^. First, a notched hydrogel samples (76 × 5 × 2 mm sample with a 38mm notch length) were mounted to a universal testing machine with a fixed initial distance between the upper and lower jaw face. Then it was subjected to tension force at the speed of 10mm/min until total fracture occurred. During this test, the sample was imaged with a 60frames/sec camera to capture the initial time point of notch crack propagation. The extension at this time point was considered as the critical distance between the upper and lower jaw face. Second, an un-notched sample with the same dimension as the notched sample was subjected to the same tension force in order to plot the force-distance curve. The area under the force-distance curve from zero to the critical distance was measured as U(h_c_) and the fracture energy (Γ) was calculated with the following equation;

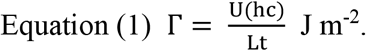

#### FTIR measurement

We prepared 3% agarose gel, PAAm gel, and agarose:PAAm hybrid gels. For the PAAm gel, 10w/v% AAm:MBA=19:1 in DI water was mixed with 0.015 w/v% APS, 0.15% TEMED and UV crosslinked with UV light (17mw/cm^2^, 25minutes). Each gel was air dried in a ventilated hood overnight. The resulting dried films were used for FTIR absorbance measurement. FTIR spectra were recorded between 4000 and 400 cm^−1^ on a Thermo Scientific™ Nicolet™ iS™ 10 FT-IR spectrometer.

#### ^13^C NMR chemical shift measurement

Nuclear magnetic resonance chemical shifts were measured by a 600MHz, high resolution NMR spectrometer (AVANCE 600, Bruker). Hybrid gel samples were prepared in D_2_O with the identical composition (agarose, acrylamide, photoinitiator) as the hybrid gels used for tensile test except MBAA which was removed to prevent excessive gel crosslinking. Agarose gel and polyacrylamide gel samples were prepared in D_2_O with 3% (w/w) and 5% (w/w), respectively. All samples contained either 0.2% Irgacure 2959 or 0.15% TEMED and 0.25% (w/v) and were exposed to UV irradiation (254nm, 14mW/cm^2^, 25min). Samples that contained agarose were boiled to completely melt the gel and was heated to 60°C during NMR measurements.

#### Performing various noncovalent hydrogel-to-hybrid hydrogel bonding

We performed the same bonding procedure of previous agarose to hybrid gel bonding except that agarose gel was replaced with alginate gel, agar gel, and gelatin gel. Exceptionally for gelatin, we crosslinked it to prevent unwanted thermal melting^33^ during UV exposure.

For agar hydrogel, mixture containing 3w/w% agar in deionized water was heated in a microwave until agar is fully dissolved. Then the mixture was poured onto glass mold and cooled at room temperature for gelation. For gelatin hydrogel, mixture containing 10 w/w% gelatin was heated in a microwave until agar is fully dissolved. Then the mixture was poured onto glass mold and cooled at room temperature for gelation. The gel was then submerged in glutaraldehyde solution (5 v/v% diluted in deionized water) overnight and then washed with deionized water. For alginate gel, pregel solution containing 2 w/w% sodium alginate was mixed to 100mM Tris-HCl buffer (pH 7.4) and dissolved by rotating the container overnight on a rotator (5rpm). The pregel solution was thoroughly degassed and mixed with a slurry of containing ionic crosslinker (20mM CaSO_4_) and crosslinking attenuator (2mM TPP), mixed until homogenous, poured into glass mold, and let to set in vacuum chamber for 1 hour. Before gelation, each pregel solution was mixed with food dye of designated color for final visualization.

Bonding each gel to hybrid gel was performed as same as agarose bonding to hybrid gel. Benzophenone treatment was omitted for negative control bonding samples. Although we could have skipped the BP treatment step by using Irgacure 2959-based hybrid instead of APS/TEMED, Irgacure 2959 made the UV-crosslinked hybrid gel slightly more brittle thus less preferred for bending tests.

Speculatively, for selecting other polypeptide-based noncovalent hydrogel candidates for bonding, one should consider ones that have i) significantly high water content and ii) moderate nonpolar amino acid content so that proper monomer/initiator diffusion occurs. One should also consider iii) UV-transparency of the hydrogel since UV must penetrate through the bulk hydrogel for proper crosslinking.

#### Preparing solids and elastomers to bond with hybrid gels

For glass or aluminum substrates, we adapted a previously reported TMSPMA-coating protocol^2^. Briefly, air plasma-activated glass, aluminum substrates were submerged in DI-based solution containing 1 wt.% TMSPMA, 0.1 v/v% acetic acid for 1-2 hours. The substrates were washed thoroughly with methanol and dried with air gun.

For PDMS substrates, PDMS film was air plasma-activated, treated with BP solution (10 w/v% in absolute ethanol), washed twice with methanol, and dried with air gun.

#### Fabrication of micropatterned intestinal model

A raster scanning microchannel (500μm x 100μm cross section) patterned agarose film (3 wt.% in DI) was obtained using a SU8 mold. The film was then attached to a hybrid film (containing 0.2% Irgacure 2959). The hybrid was pat-dried before attachment. The attachment was askew to leave a portion of the hybrid exposed. The exposed part of the hybrid was covered with aluminum foil before UV exposure (17mW/cm^2^, 10 min). The structure was folded into a cylinder, anchored so that the UV-unexposed portion of hybrid remains attached to the dangling agarose end. The structure was then exposed to additional UV light (17mW/cm^2^, 10 min). A graphical description is given in supplementary figure 7.

#### Fabrication of microfluidic DNA electrophoresis sample retriever

Two agarose gels (1.5 wt.% in 0.5x TBE, contained 10x SYBR safe), where one gel contained DNA sample inlet, was attached by an intermediate, 1mm-thick hybrid gel film pre-patterned with an internal channel. The construct was exposed to UV light (17mW/cm^2^, 40min) while the agarose gel partitions were protected with aluminum foil. TMSPMA-coated glass pipette was coated with agarose-AAm pregel solution, inserted into the inlet and outlet of the hybrid channel. The construct was again exposed to UV light (17mW/cm^2^, 40min) while the agarose gel partitions were protected with aluminum foil. We connected the outlet glass pipette with a 1x TE solution-containing syringe by tygon tubing. We injected TE solution so that the hybrid channel is filled with TE solution without bubble. The construct was then submerged in 0.5x TBE (running buffer). After 2.5uL of 100bp ladder sample was injected into the sample inlet, electrophoresis was performed (approximately 6V/cm). We manually observed the size separated DNA ladders so that when the target band (1.4kbp) fully enters the channel, we could retrieve the sample by withdrawing the syringe. Retrieval yield was analyzed by measuring the amount of dsDNA with Qubit dsDNA (broad range mode, Invitrogen) and comparing with the input amount. Conventional agarose gel electrophoresis was used to measure purity; examining contamination from other DNA bands.

## Acknowledgments

This research was supported by Global Research Development Center Program through the National Research Foundation of Korea(NRF) funded by the Ministry of Science and ICT(MSIT) (2015K1A4A3047345)

## Author contributions

S.W.B., H.S.K., and J.-Y.S. designed and conducted experiments. S.W.B. wrote the manuscript. Y.-W.K. performed tensile tests. H.J.B. performed SEM imaging. S.W.S., H.Y., and S.D.K. provided biochemical reagents, tools, and advices regarding experiments. J.-Y.S. and S.K. overviewed writing the manuscript.

### Additional information

**Supplementary information** accompanies this paper at http://www.nature.com/naturecommunications

## Competing financial interests

The authors declare no competing financial interests.

**Reprints and permission** information is available online at http://npg.nature.com/reprintsandpermissions/www.nature.com/reprints.

