## supplementary figures for "Robust hydrogel-integrated microsystems enabled by enhanced interfacial bonding strength"

### Supplementary Materials

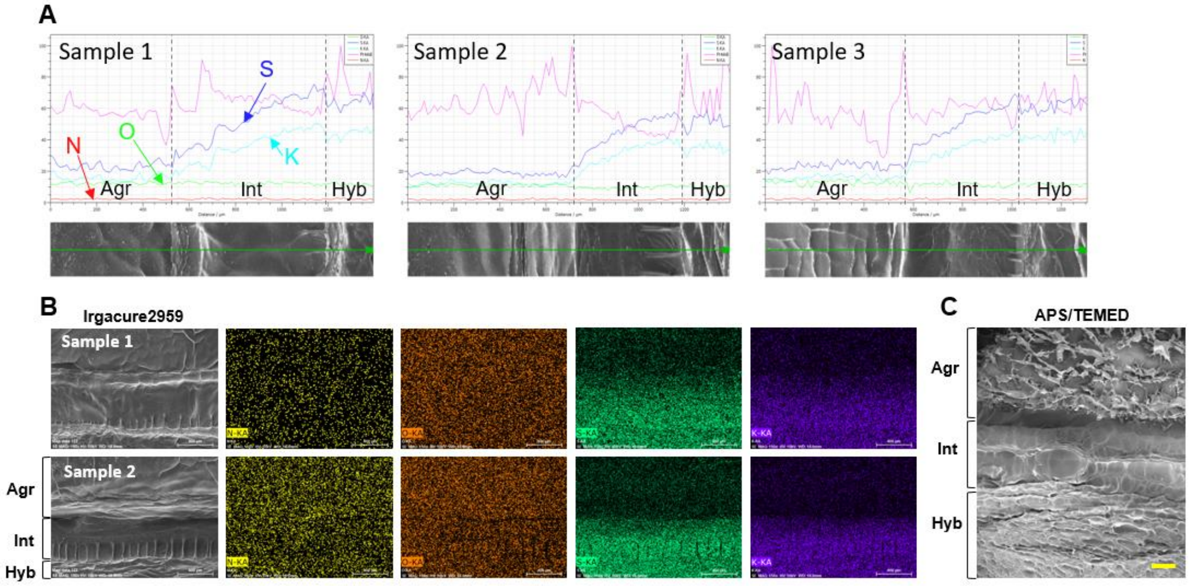


**Figure S1. Validation of bonding mechanism using SEM/EDS measurement. A,B.** EDS line-scan mode result from 3 different samples (A) and area profiling from 2 samples (B). SEM image and EDS profile of agarose:hybrid interface fabricated by Irgacure2959 photo-initiation. Sulfur and potassium signal is saturated at the Hybrid (Hyb) region, decreases along the interface (Int), and remains base level throughout the agarose (Agr) region. Whereas, nitrogen and oxygen signal remains base level across the three layers. **C**. SEM image of the interface with a different photo-initiation method using APS and TEMED. Scale bar is 100um.

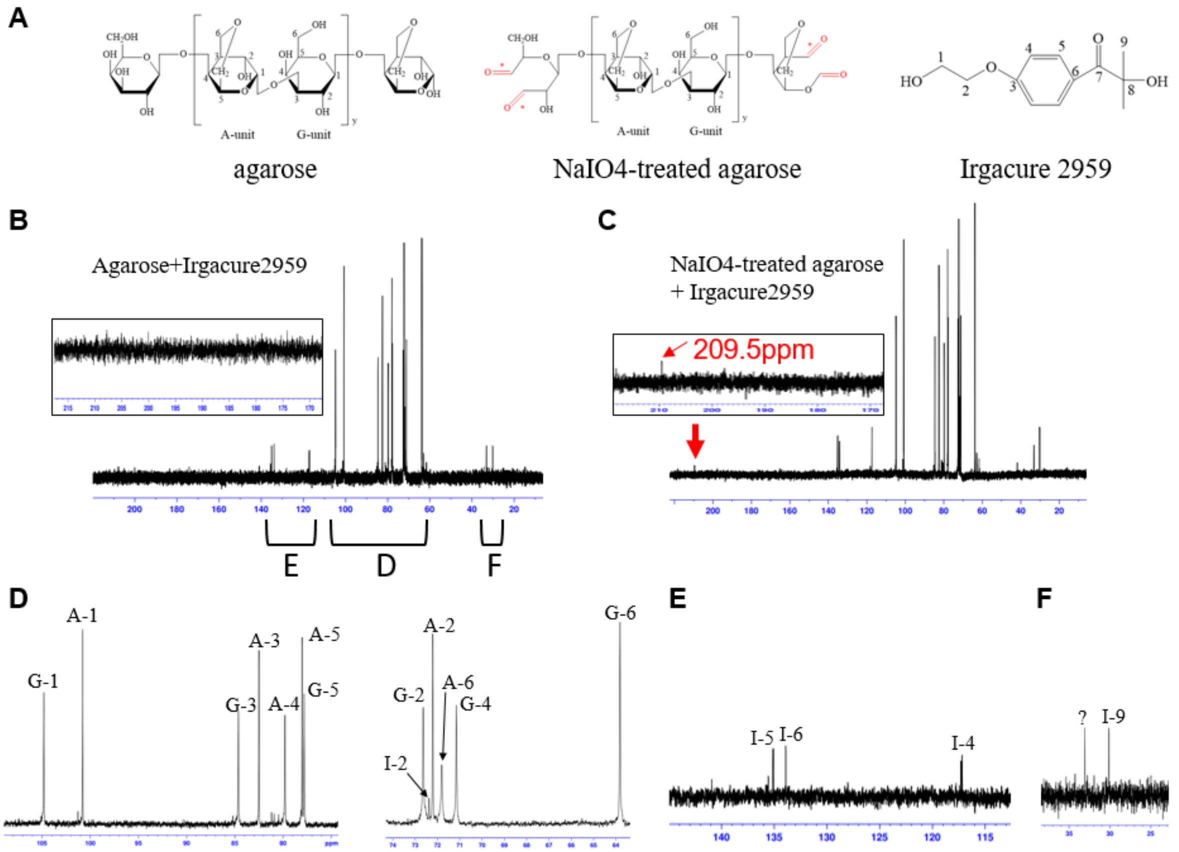


**Figure S2. Verification of aldehyde-activation of agarose by sodium periodate treatment. A.** Predicted chemical structures of agarose, sodium-periodate-treated agarose, and Irgacure 2959. **B,C.** $^{13}$C-NMR spectrum of native agarose (B), and sodium-periodate-treated agarose (C). Inset of B and C shows that only sodium-periodate-treated agarose show shift at 209.5 ppm (red arrow), which corresponds to the carbonyl group of aldehyde. D-F. Magnified image of the areas indicated in (B). The chemical shifts corresponding to agarose A-unit, agarose G-unit, and Iragcure 2959 are marked as A-x, G-x, I-x, respectively.

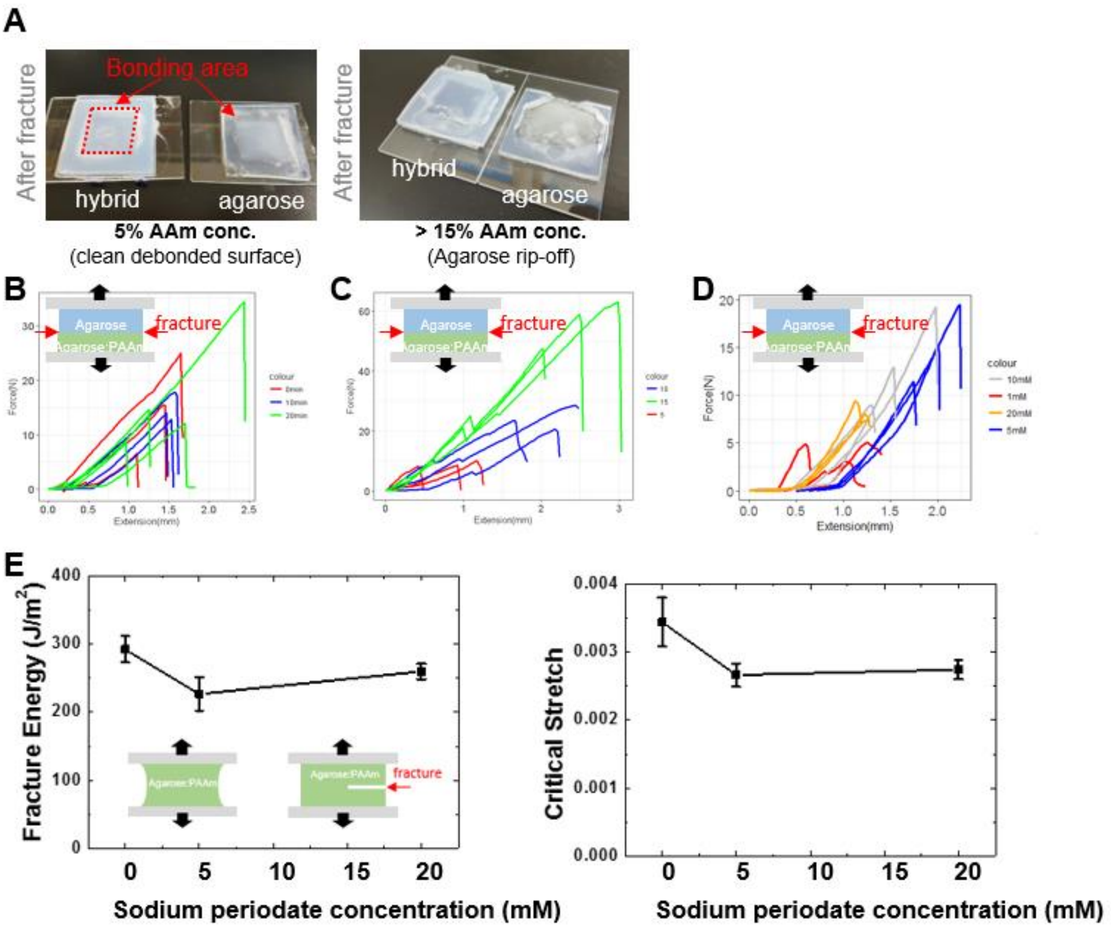


**Figure S3. Measuring bonding strength using flatwise tensile test. A**. Examples of de-bonded samples after tensile test. **B-D.** Raw data of flatwise tensile tests for figure2c,d, and figure S4c, respectively. **E**. Result of fracture energy and critical stretch measurement of hybrid hydrogels. Hybrid hydrogels contained 5 w/v% AAm:MBAA=19:1, 3 w/w% aldehyde-activated agarose. Aldehyde activation was performed with either 0, 5, 20mM sodium periodate for 3 hours.

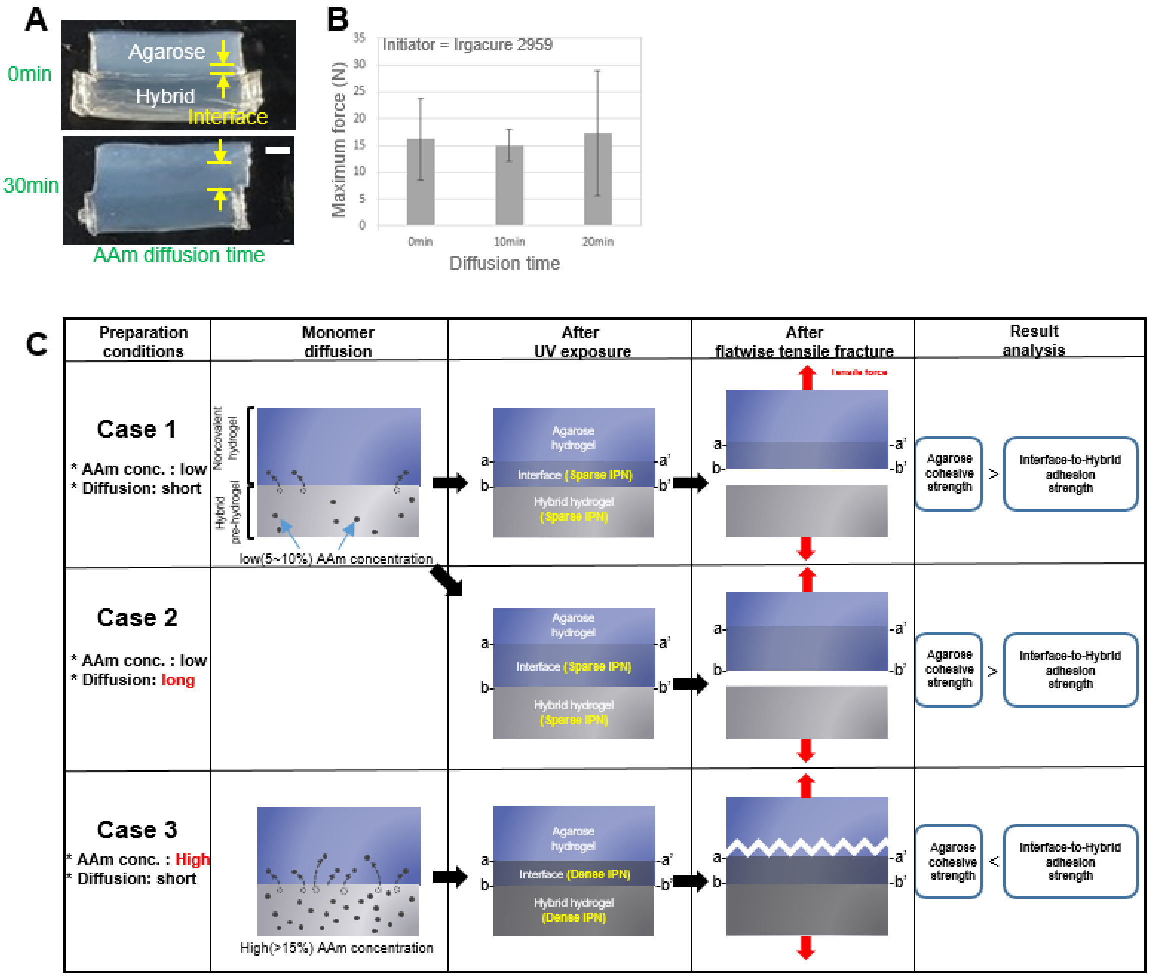


**Figure S4. Varying monomer diffusion time has no significant effect on the variation of bonding strength. A, B.** Cross-section (A) and maximum force measurement (B) of the bonded agarose gel-hybrid hydrogel structures that were incubated at ambient condition for monomer diffusion for different duration before UV exposure. **C.** Graphical explanation of the phenomenon. For Case 1 and 2, When the initial hybrid gel contained low (or moderate) AAm concentration, the resulting bonding strength didn't vary much with different "monomer diffusion time" because the debonding plane formed consistently at b--b' and not within the interface volume (between a--a' and b--b'). We think that the debonding plane formed at b--b' because the agarose's matrix cohesive strength exceeded the strength of interface-to-hybrid adhesion. Therefore, the debonding plane (the plane of weakest strength) wouldn't change by varying the "monomer diffusion time" (or the interface thickness). For Case 3, as we shown in A, when hybrid's AAm concentration was high (>15%), the interface-to-hybrid adhesion strength exceeded the agarose's matrix cohesive strength, and the resulting debonding plane occurred within the agarose gel matrix. This led to agarose gel tear off (as shown in A, right-side figure). In this case, even if we had vary the diffusion time (interface thickness) the bonding strength we measure would just have been the agarose gel's cohesive force, not the bonding strength between agarose and hybrid gel. Hybrid gel contained 3% agarose activated with 24 hour treatment of 100mM $NaIO_4$, 10% (w/v) AAm:MBA=19:1, 0.2% Irgacure 2959. Scale bar is 2.5mm (A),

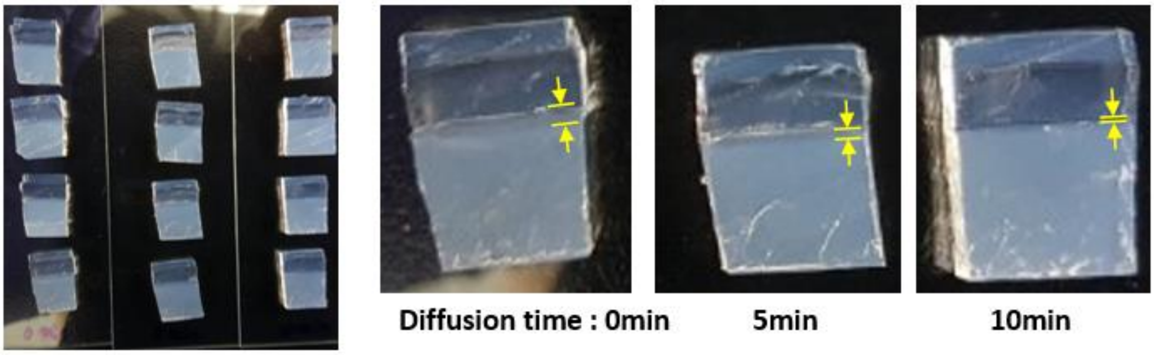


**Figure S5. Observation of decreasing interfacial layer thickness with increased monomer diffusion time when using APS/TEMED-based photoiniation.** When hybrid gel was fabricated using APS/TEMED as photoinitiator and catalyst, the interfacial layer thickness decreased when passive monomer diffusion time was increased. This was due to exposure to oxygen over time.

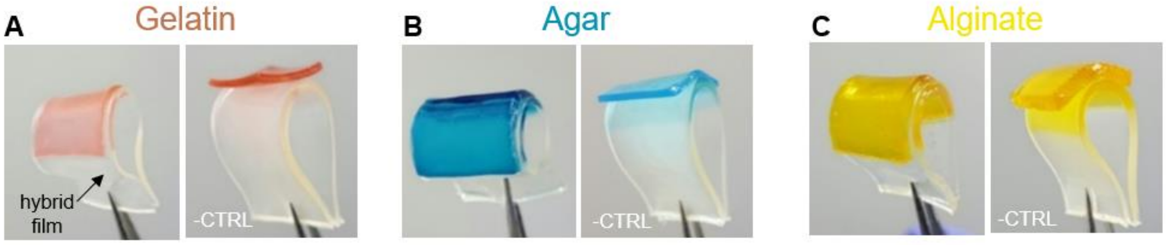


**Figure S6. Bonding method is adaptable to other noncovalent hydrogels.** Bonding Hybrid hydrogel with gelatin (A), agar (B), and alginate (C) hydrogel. The hybrid hydrogel film was composed of agarose:AAm and Irgacure 2959 initiator. –CTRL indicates negative controls showing bonding failure. For negative control, hybrid gel was partially UV-crosslinked with PDMS film before loading the noncovalent hydrogel and subsequent UV exposure.

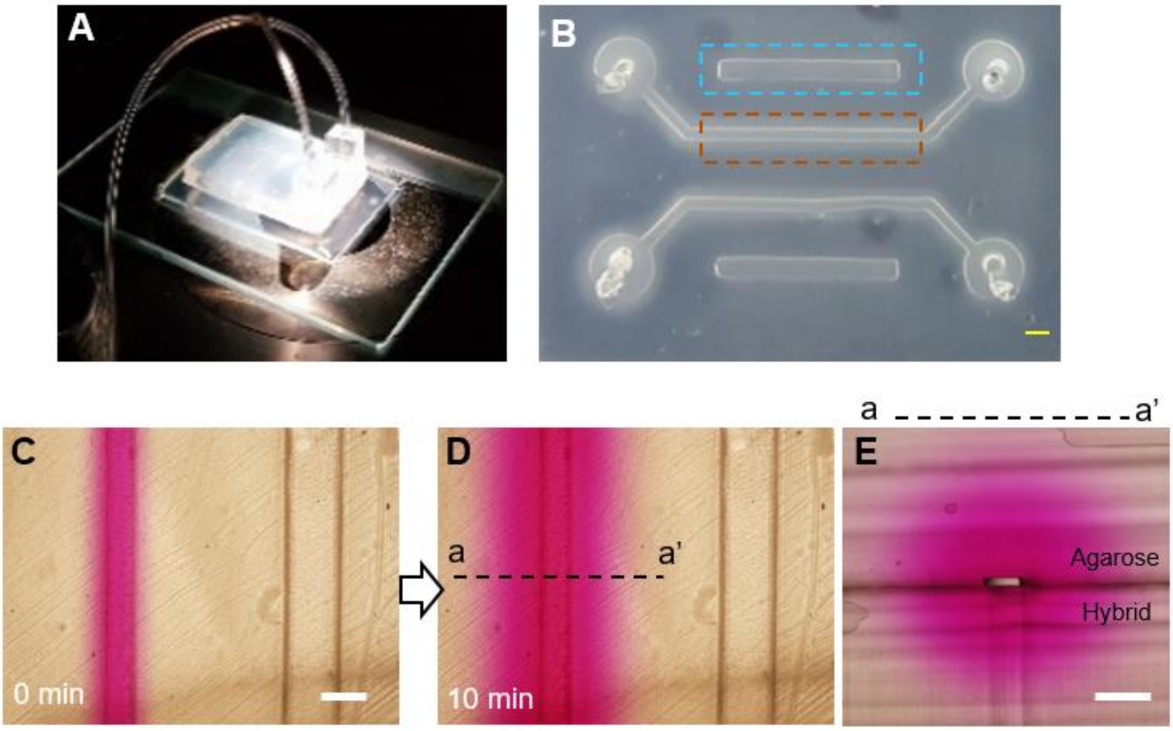


**Figure S7. Diffusive microfluidic test imaged at bright field mode.** A mechanical anchorage-free diffusive microfluidics demonstration (A). (B) Rhodamine B was injected into a microchannel (indicated with dotted red box) patterned between the agarose to hybrid bonding interface. After 10 minutes of dye injection (C, D), cross section was imaged (E). Scale bars are 500um.

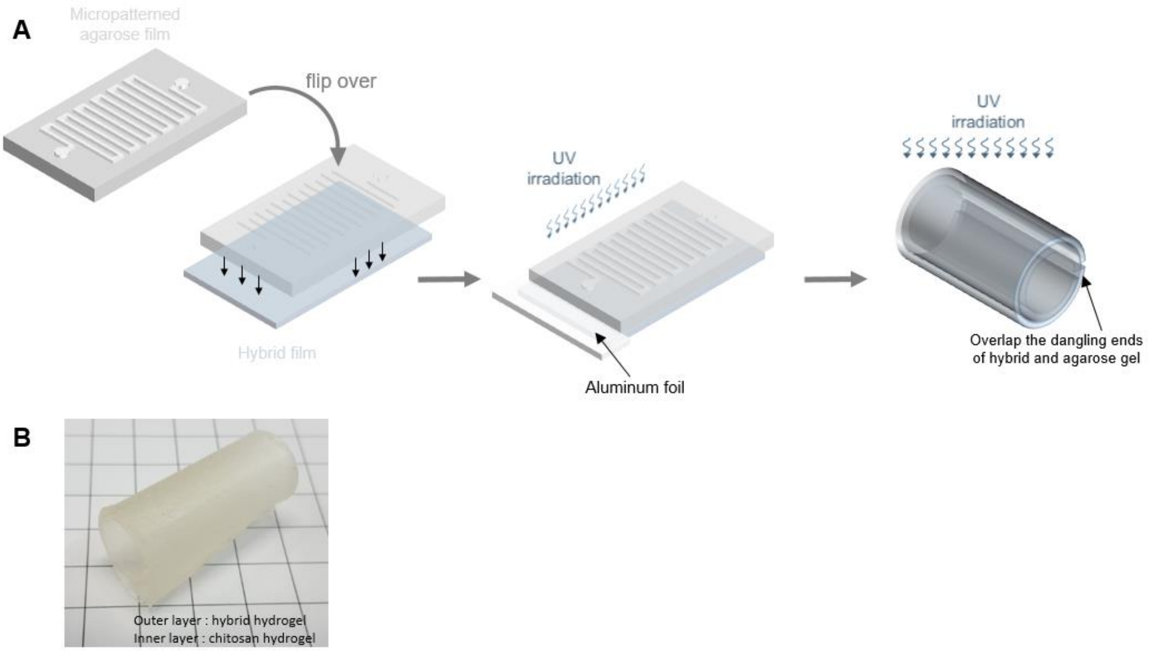


**Figure S8. Fabrication process and example of hollow intestinal organ model.**
(A) Intestine-like model fabrication process. (B) Example fabrication product containing chitosan hydrogel inner layer and agarose/acrylamide hybrid hydrogel outer layer.

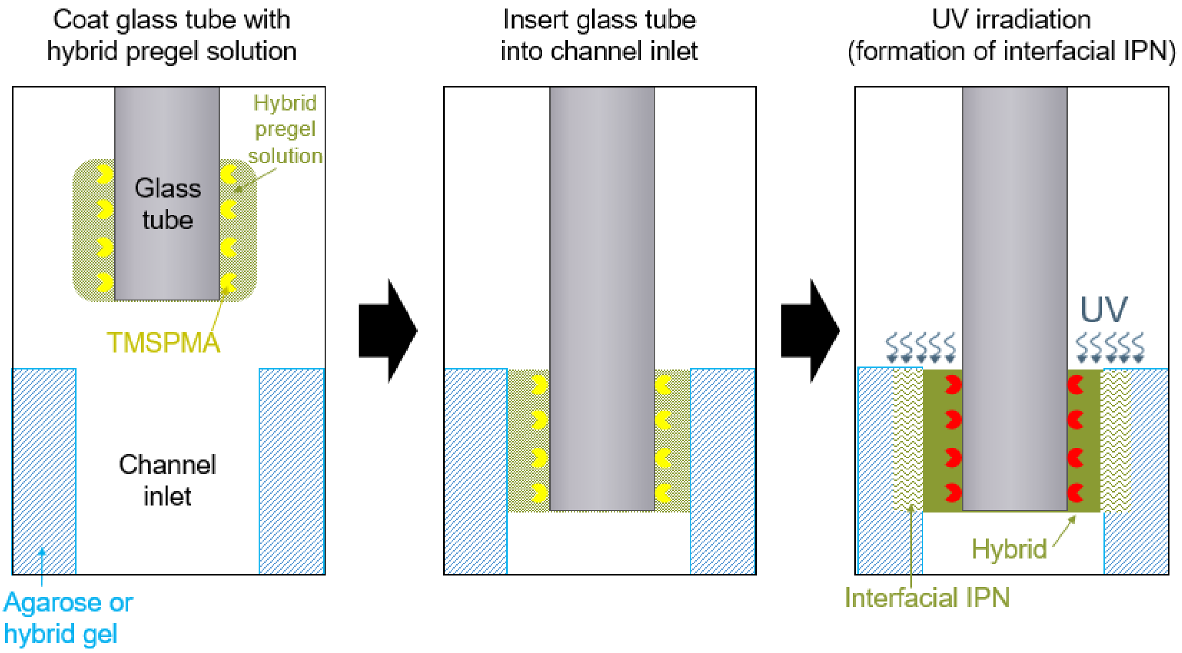


**Figure S9. Glass tube to hydrogel bonding method.** A glass pipette coated with hybrid pregel-solution is inserted into the inlet and outlet of the channel and fixed by UV curing. UV curing creates interfacial IPN between the glass-coated hybrid and the wall of the hydrogel channel as well as chemical bond between glass and hybrid gel.

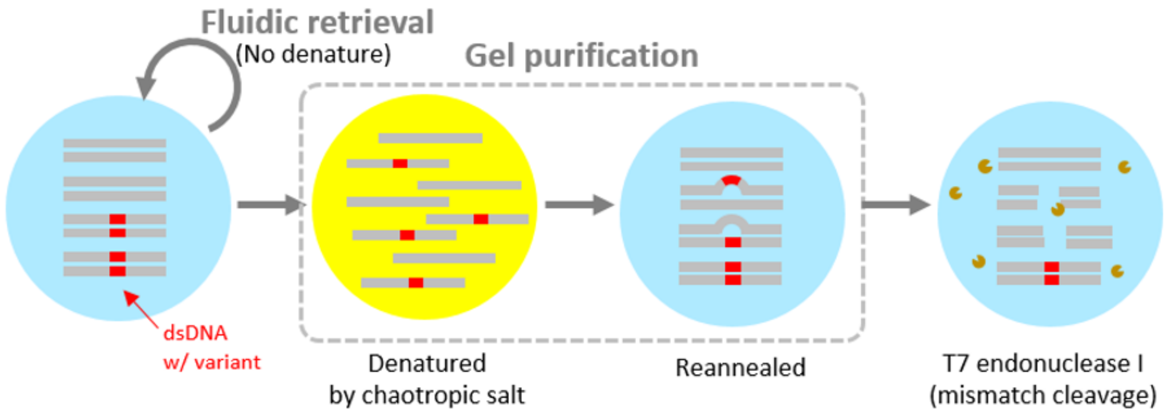

**Figure S10. Schematic showing the effect of the retrieval method on the structure of retrieved double strand DNA (dsDNA) sample.** The original input sample (left) consists of dsDNA that have identical sequences at most part (gray) but a proportion of the sample contains DNA variants (red) such as polymorphism or mutation or amplification error. When the samples are purified with commercial gel purification, the reagent contains chaotropic salts which are used to denature agarose gels by interfering hydrogen bond. This salt however also denatures dsDNA into single-strand DNA (ssDNA). After salt is removed and DNA samples are re-suspended in low-salt water, the ssDNA strands reanneal into dsDNA. During this reannealing step, the ssDNA that contain variant could erroneously anneal with non-variant-containing ssDNA and form a mismatched dsDNA. This mismatch results in a bulge in the dsDNA with can be detected by T7 endonuclease I enzyme and cleaved. The result is fragmented dsDNAs (right). Compared to this, fluidic retrieval method doesn’t involve any gel-melting step, and thus dsDNA samples retain their original structure.
